# Time course of the physiological stress response to an acute stressor and its associations with the primacy and recency effect of the serial position curve

**DOI:** 10.1101/568352

**Authors:** Linda Becker, Nicolas Rohleder

## Abstract

Whether stress affects memory depends on which stress pathway becomes activated and which specific memory system is involved. The activation of the sympathetic nervous system (SNS), leads to a release of catecholamines. The activation of the hypothalamic-pituitary-adrenal (HPA) axis, leads to a release of glucocorticoids. In thus study, it was investigated whether SNS and/or HPA axis activation are associated with long-term memory (LTM) and/or working memory (WM) performance in humans. Thirty-three participants underwent the socially evaluated cold-pressor test. Salivary alpha-amylase (sAA) was used as a marker for the activation of the SNS and cortisol as marker for HPA axis activation. Memory was assessed by means of word lists with 15 words each. The primacy effect (i.e., the correctly recalled words from the beginning of the lists) of the serial position curve was considered as indicator for LTM. The recency effect (i.e., the correctly recalled words from the end of the lists) were used as estimator for WM performance. In sAA responders, the recency effect and, therefore, WM performance increased immediately after the stressor. This was not found in sAA non-responders. In cortisol responders, the primacy effect and, thus, LTM performance decreased 20 minutes after the stressor. No change in LTM performance was found in cortisol non-responders. Our study supports the assumptions that 1) SNS activation is associated with WM processes via stimulation of the prefrontal cortex, and 2) HPA axis activation is associated with LTM processes through interactions with the hippocampus.

## Introduction

Cognitive functions – especially memory – are not entirely independent of peripheral physiological processes. Some peripherally transmitted molecules (e.g., some hormones) can pass the blood-brain barrier (BBB) and can, therefore, affect neural activity directly. Other substances indeed cannot pass the BBB but can still affect neural activity through indirect feedback loops by activating brain networks, which lead to the release of neuro modulators in brain regions involved in cognitive processing (e.g., 1–3).

One prominent candidate which can trigger such effects is stress. Stress can be defined as “*an actual or anticipated disruption of homeostasis or an anticipated threat to well-being*” (4, p. 397). The acute stress response is dominated by two pathways (e.g., 5; 6, Fig 1a). The first, which starts immediately after the onset of the stressor is the activation of the sympathetic nervous system (SNS). This leads to the release of the catecholamines adrenaline and noradrenaline which both cannot pass the BBB but can affect cognitive processing through indirect pathways (1). The peripherally transmitted catecholamines can activate the locus coeruleus in the brainstem which stimulates the release of noradrenaline and dopamine in the prefrontal cortex (PFC) via the ventral tegmental area (7; 1, 8). The PFC is involved in a variety of higher order cognitive functions, e.g., in working memory (WM) processes which are mainly controlled by noradrenaline and dopamine (9).

**Fig 1.**
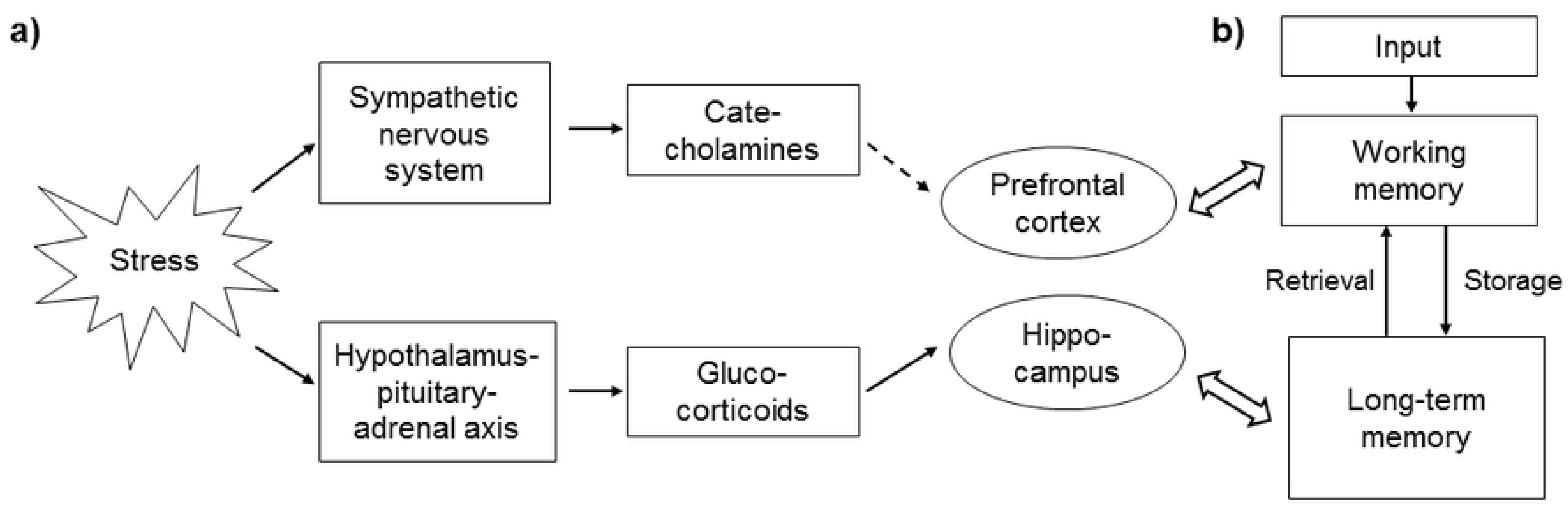
a) Stress pathways that are activated after an acute stress situation, affected brain regions through indirect (dashed arrow) or direct pathways, and b) a simplified memory model.

The second stress response, which peaks with a short delay of a few minutes after the onset of the stressor, is the activation of the hypothalamic-pituitary adrenal (HPA) axis. This leads to the release of glucocorticoids (i.e., cortisol in humans or corticosterone in rodents) from the adrenal cortex. After threatening socially-evaluative stressors (e.g., the Trier Social Stress Test; 10), HPA axis response peaks approximately 20 minutes after the end of the stressor (e.g., 11). The stress hormone cortisol can pass the BBB and can, therefore, directly affect neural processing (12). Cortisol binds to two different receptors in the brain (13, 14). The first, the mineralocorticoid receptor (MR, or type 1 receptor) can be found within the hippocampus and the prefrontal cortex (15). The second, the glucocorticoid receptor (GR, or type 2 receptor) is widely distributed in different brain areas. Which cognitive processes (i.e., which memory functions) are affected after cortisol release depends on which receptors and, therefore, in which brain area, cortisol binds to (16). Both receptor types have different affinity for cortisol (13). The MRs have high affinity and are, therefore, usually occupied at basal cortisol concentrations. The GRs have a lower affinity for cortisol and are, thus, in many cases not occupied unless cortisol levels are increased. The brain structure in which both MRs and GRs are localized is the hippocampus which is also mainly involved in long-term memory (LTM) processes. Therefore, there has been a long research history in the evaluation of the effects of cortisol binding on GRs in the hippocampus and its associations with LTM processes (e.g., 17; 18, 19).

A classical model which involves both WM and LTM, was proposed by 20 (20). According to this model, any new information first enters – after it has passed the so-called sensory register – WM. After this, the information is either forgotten or it is stored in LTM from where it can be retrieved at later time points (Fig 1b). One easy way to assess both WM and LTM within one experiment is to investigate the so-called serial position effects (21). A typical experimental procedure is to present word lists of approximately 15 words to the participants and to let them recall as many words as they can remember immediately after the last word was presented. The classical observation is that words from the beginning as well as from the end of the word list can be better recalled than words from the middle of the list. The first effect is called the primacy effect (PE) which is associated with LTM processes. The second is called recency effect (RE) and it is associated with WM (22). The effects of acute stress on the serial position effects have not been investigated so far.

The aims of the present study were to investigate whether SNS activation is associated with WM and whether HPA axis activation is associated with LTM performance in humans after an acute stressor. As measures of the functioning of the memory systems, the PE and the RE of the serial position curve were examined. The hypotheses were that 1) SNS activation would start immediately after the stressor and would be related with the RE and, thus, with WM and 2) that HPA axis activation would peak with a time delay of approximately 20 minutes and would be associated with the PE and, therefore, with LTM performance. It was assumed that these effects will only be found in participants who show indeed a SNS or HPA axis response, respectively (the so-called responders). Furthermore, it was hypothesized that such effects will not be found in the non-responder groups.

## Materials and methods

### Participants

Thirty-three healthy, German-speaking adults participated (mean age: 24.0 ± 5.7 years; eight male; BMI = 22.2 ± 2.8 kg/m^2^). None of them reported endocrinological, neurological, or psychological diseases. All participants gave their written and informed consent. The study was carried out in accordance with the Code of Ethics of the World Medical Association (Declaration of Helsinki) and was approved by the local ethics committee of the Friedrich-Alexander University Erlangen-Nuremberg (protocol # 6_18 B).

### Experimental procedure

The time course of the experiment is shown in Fig 2. The whole session – including instructions – lasted 60 minutes. For memory assessment, participants were presented three word lists with 15 words each with inter-stimulus intervals of one second. The words were simple neutral words with a short pronunciation time (e.g., the German words for ‘dog’, ‘coffee’, ‘bus’, or ‘door’). After the presentation of each list, the participants were asked to immediately recall as many words as they had remembered. As measure for LTM performance, the PE was used which was defined as the sum of correctly recalled words from the first three words of the lists. Accordingly, the RE, which was used as a measure for WM performance, was defined as the sum of correctly recalled words from the last three words of the lists. Memory testing was repeated three times throughout the experimental session with three lists each time. The order of the word lists was counterbalanced between the participants and between the memory assessment time points.

**Fig 2.**
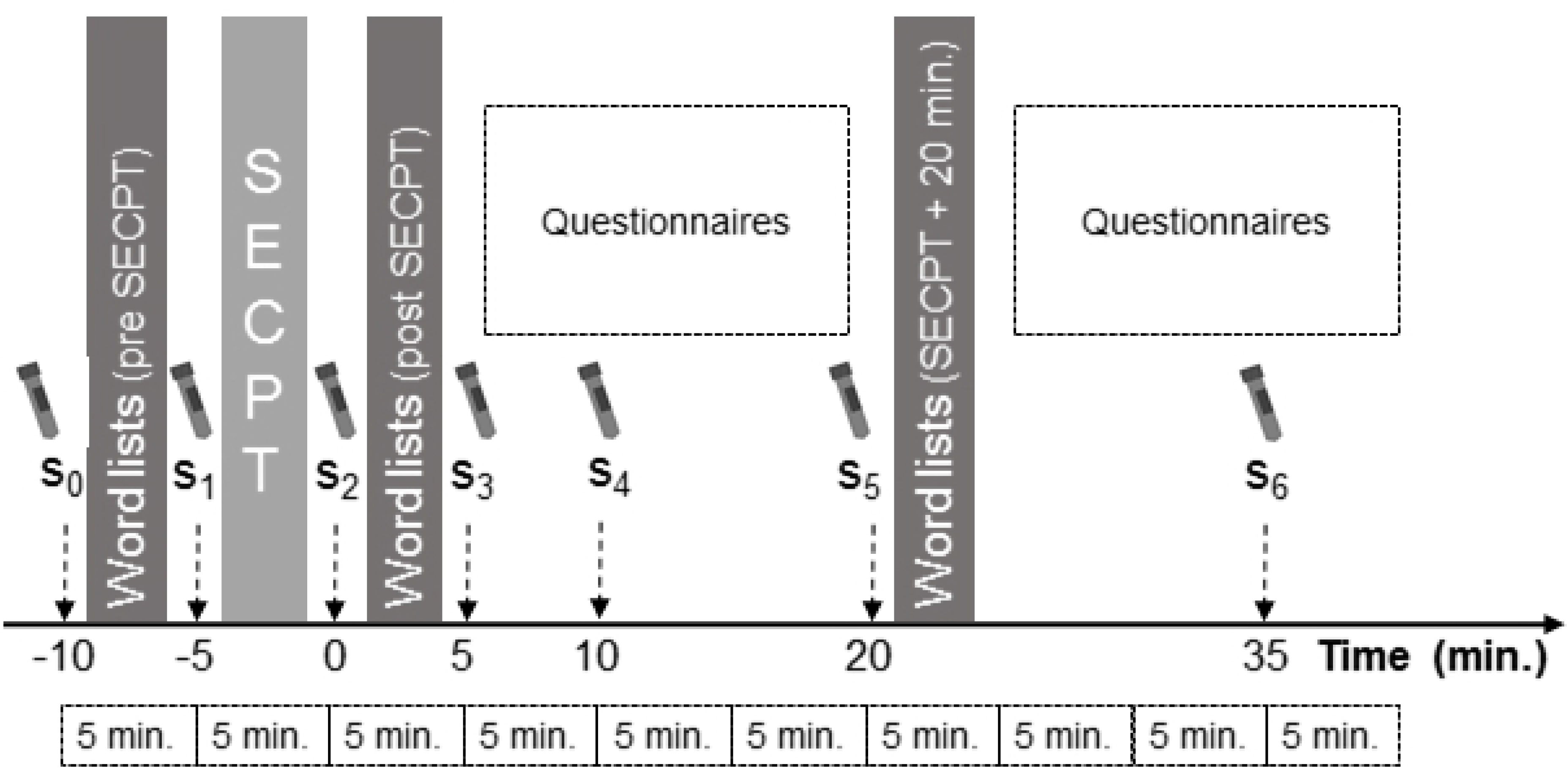
Time course of the experimental procedure. Stress was induced by means of the socially evaluated cold-pressor test (SECPT).

Stress was induced by means of the socially evaluated cold-pressor test (SECPT, 23) in groups of two participants. The participants stood in front of a table on which transparent boxes filled with ice water were placed. Participants were instructed to immerse their hands in the ice water as long as possible for up to three minutes. Mean immersion time was 2:20 ± 1:06 min (min. 0:18, max: 3:00). The hand of each participant was directly opposite of the hand of the other person with the aim to introduce a competitive situation. Remaining time was displayed on a large-display digital clock that was visible for both participants. An auditory countdown announced the last five seconds. Therefore, our protocol slightly differed from that reported by Schwabe and colleagues (2008) and by Minkley and colleagues (2014) who introduced the first group version of the SECPT. One experimenter who wore a medical uniform was present during the SECPT and was instructed to behave distanced and to keep a neutral mimic.

Salivary alpha-amylase (sAA) and salivary cortisol were used as measures of SNS and HPA axis activity (24–27). Saliva was collected by means of salivettes (Sarstedt, Nümbrecht, Germany) at seven time points during the experimental session. The first saliva sample (s_0_) was collected immediately prior to the presentation of the first word list. The following samples were collected immediately prior (s_1_) and immediately after (s_2_) the SECPT. The following four samples were collected five (s_3_), ten (s_4_), 20 (s_5_), and 35 (s_6_) minutes after the end of the SECPT. The participants were instructed not to eat, drink (except water), smoke, or brush their teeth two hours before the start of the experimental session. Additionally, subjective stress perception was rated on a ten-point Likert-scale with the anchors *“not stressed at al;*” and *“totally stressed*” during saliva collection.

Furthermore, some demographic and psychological variables were collected by means of questionnaires during waiting time between the saliva samples (when no memory tests were performed). Demographic variables that were assessed were sex, age, weight, and height. The amount of regular physical activity was measured by means of the short form of the International Physical Activity Questionnaire (IPAQ; 28; 29). Chronic stress was assessed by means of the screening scale of the Trier Inventory of Chronic Stress (TICS-SSCS; 30) and the Perceived Stress Scale (PSS; 31). Additionally, burnout and depression were measured by means of the Maslach Burnout Inventory (32) and the German version of the depression scale from the Center for Epidemiological Studies (CES-D, 33, 34).

### Sample processing

Saliva samples were stored at −30 °C after collection for later analyses. After the study was completed, samples were sent to Dresden LabService GmbH (Dresden, Germany) where they were analyzed by means of high performance liquid chromatography.

### Sample analysis

For statistical analyses, IBM SPSS Statistics (version 25) was used. For evaluation of the memory test, the number of recalled words for each position (1 to 15) was summed for the three word lists at each the three measurement time points. First, only the memory tests were analyzed to ensure that primacy and recency effects were actually found.

Therefore, an analysis of variance for repeated measurement (rmANOVA) with the within-subject factors ‘position’ (‘1’, …, ‘15’) and ‘time point (‘pre SECPT’, ‘post SECPT’, and ‘SECPT + 20 min.’) was calculated. The number of correctly recalled words was averaged over the percentiles (1^st^ to 3^rd^, 4^th^ to 6^th^, …, 13^th^ to 15^th^ word position; P_1_, …, P_5_) to make the following post-hoc analysis easier to interpret. Only the percentiles were used for further statistical analysis. If necessary, sphericity violations (determined by Mauchly’s test of sphericity; 35) were corrected by adjusting the degrees of freedom with the procedure by 36. As post-hoc tests, *t*-tests with adjusted alpha levels according to the Bonferroni correction were calculated. Partial eta-squares (η_p_^2^) for ANOVAs and Cohen’s *d* for *t*-tests are reported for effect sizes. If necessary, Cohen’s *d* was corrected according to the method that was proposed by 37 (37). For further analysis (after the occurrence of an PE and RE was revealed), P_1_ was considered as a measure of the primacy effect and, therefore, for long-term memory, and P_5_ was considered as a measure for the recency effect and, thus, for working memory performance. To test whether the PE and the RE differed between the three time points (‘pre SECPT’, ‘post SECPT’, and ‘SECPT + 20 min.’) and whether they were, therefore, related to the stress induction, a further rmANOVA with the factors ‘time point’ and ‘memory effect’ (‘PE’ and ‘RE’) was calculated.

Because of positive skewness, cortisol levels were transformed by means of the natural logarithm prior to further statistical analysis. Participants were classified as responders or non-responders, separately for sAA and cortisol. Participants with an increase of more than 10 percent between s1 and s2 for sAA and between s1 and s5 for cortisol were classified as responders. Further rmANOVAS with the within-subject factors ‘memory type’ and ‘time point’ and the between-subject factor ‘respondence’ were calculated. If necessary, post-hoc rmANOVAS with the factors ‘time point’ and ‘respondence’ were calculated, separately for the PE and RE.

## Results

### Stress effects on memory

A main effect of the factor position (F_(6.011,192.35)_ = 24.80, *p* < .001, η_p_^2^ = .44), a main effect of time point (*F*_(2, 64)_ = 4.08, *p* = .022, η_p_^2^ = .11), and an interaction position * time (*F*_(14.17, 453.42)_ = 4.83, *p* < .001, η_p_^2^ = .13) were found (Fig 3a-c). A further rmANOVA, in which the percentiles P_1_, P_3_, and P_5_ were compared, revealed a significant main effect of the factor time point (*F*_(1.61, 51.76)_ = 58.0, *p* < .001, η_p_^2^ = .64), a marginally significant main effect of the factor percentile (*F*_(2, 64)_ = 2.84, *p* = .066, η_p_^2^ = .08), and a significant interaction time point * percentile (*F*_(4, 128)_ = 13.19, *p* < .001, η_p_^2^ = .29). Post-hoc t-tests showed that both the primacy effects (i.e., P_1_ > P_3_) and recency effects (i.e., P_5_ > P_3_) were found for all three time points (all *p* < .001) in this sample.

**Fig 3.**
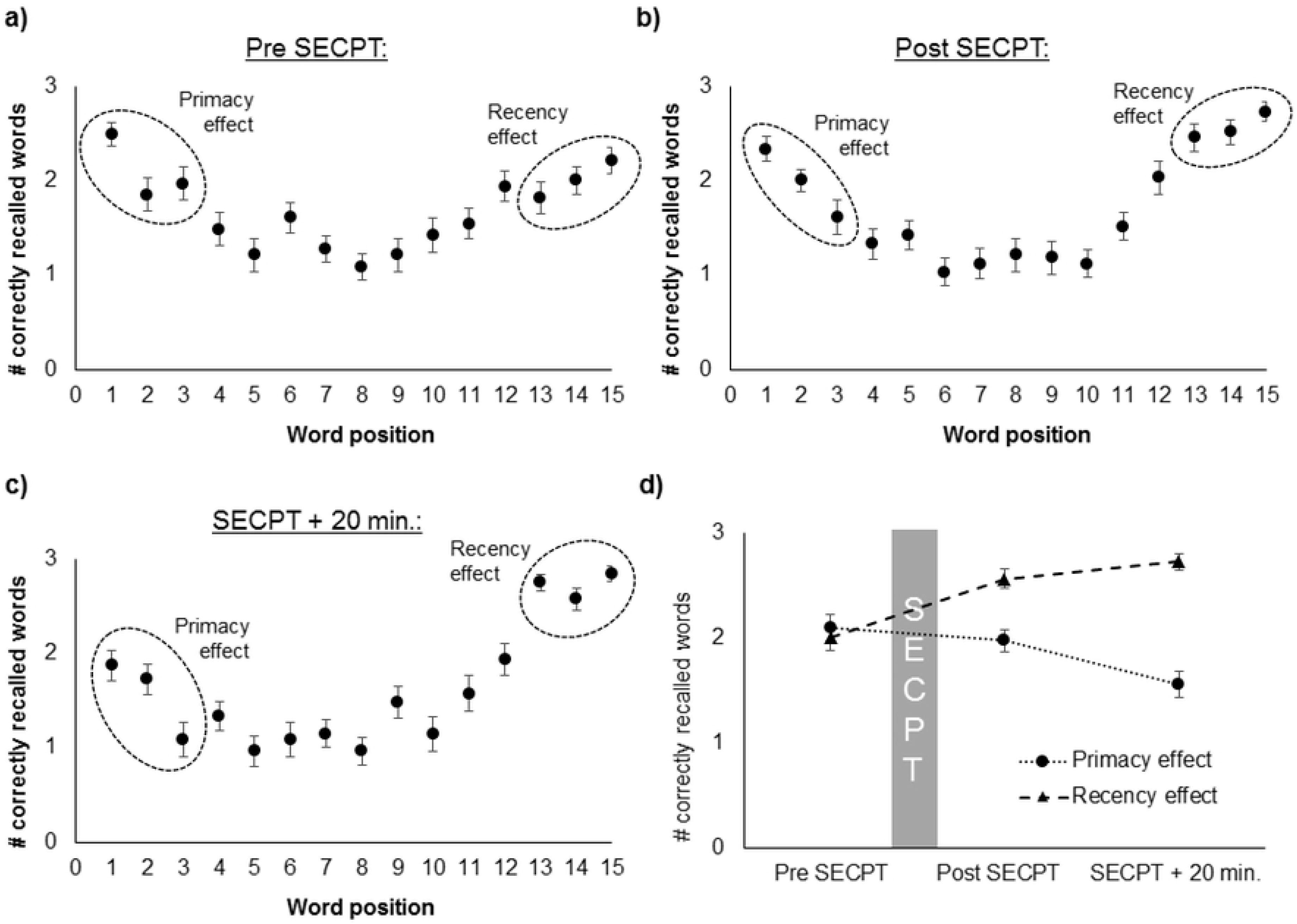
Number of correctly recalled words in dependence of the word position, a) before the SECPT, b) immediately after SECPT, c) 20 minutes after the SECPT, and d) strength of the primacy and recency effect in dependence of the measurement time-point.

For further analysis, the first (*P_1_*) and the last (*P_5_*) three words were used as measures for the PE and RE. A further rmANOVA revealed main effects of the factors time point (*F*_(2, 64)_ = 5.76, *p* = .005, η_p_^2^ = .15) and memory effect (F_(1, 32)_ = 17.15, *p* < .001, η_p_^2^ = .35) and an interaction time point * memory effect (*F*_(1.67, 53.4)_ = 20.76, *p* < .001, η_p_^2^ = .39, Fig 3d). Post-hoc t-tests showed that the strength of the PE and RE was the same before the SECPT and different after the SECPT (i.e., the RE was stronger than the PE; both *p* < .001). Immediately after SECPT, the PE was the same as before (*p* = .236). The PE was significantly weaker 20 minutes after the SECPT than immediately after it (*t*_(32)_ = 3.65, *p* = .001, *d* = −0.71). Therefore, LTM performance did not differ immediately after SECPT, but decreased 20 minutes after the stress induction. The RE was significantly stronger immediately after the SECPT than before (pre-post: *t*_(32)_ = – 5.43, *p* < .001, *d* = 0.86), but did not change further in the following 20 minutes (*p* = .122). Therefore, WM performance increased immediately after the SECPT, but did not change afterwards.

### Subjective stress perception

Subjective stress perception significantly differed between the seven time points (*F*_(3.68, 117.75)_ = 13.56, *p* < .001, η_p_^2^ = .30). Post-hoc t-tests revealed that subjective stress perception was higher immediately after the SECPT (s_2_) than before it (s_1_-s_2_: t(_32_) = −2.97, *p* = .006, *d* = 0.56) and further declined until 20 minutes after the SECPT (s_5_, all *p* < . 230, Fig 4a).

**Fig 4.**
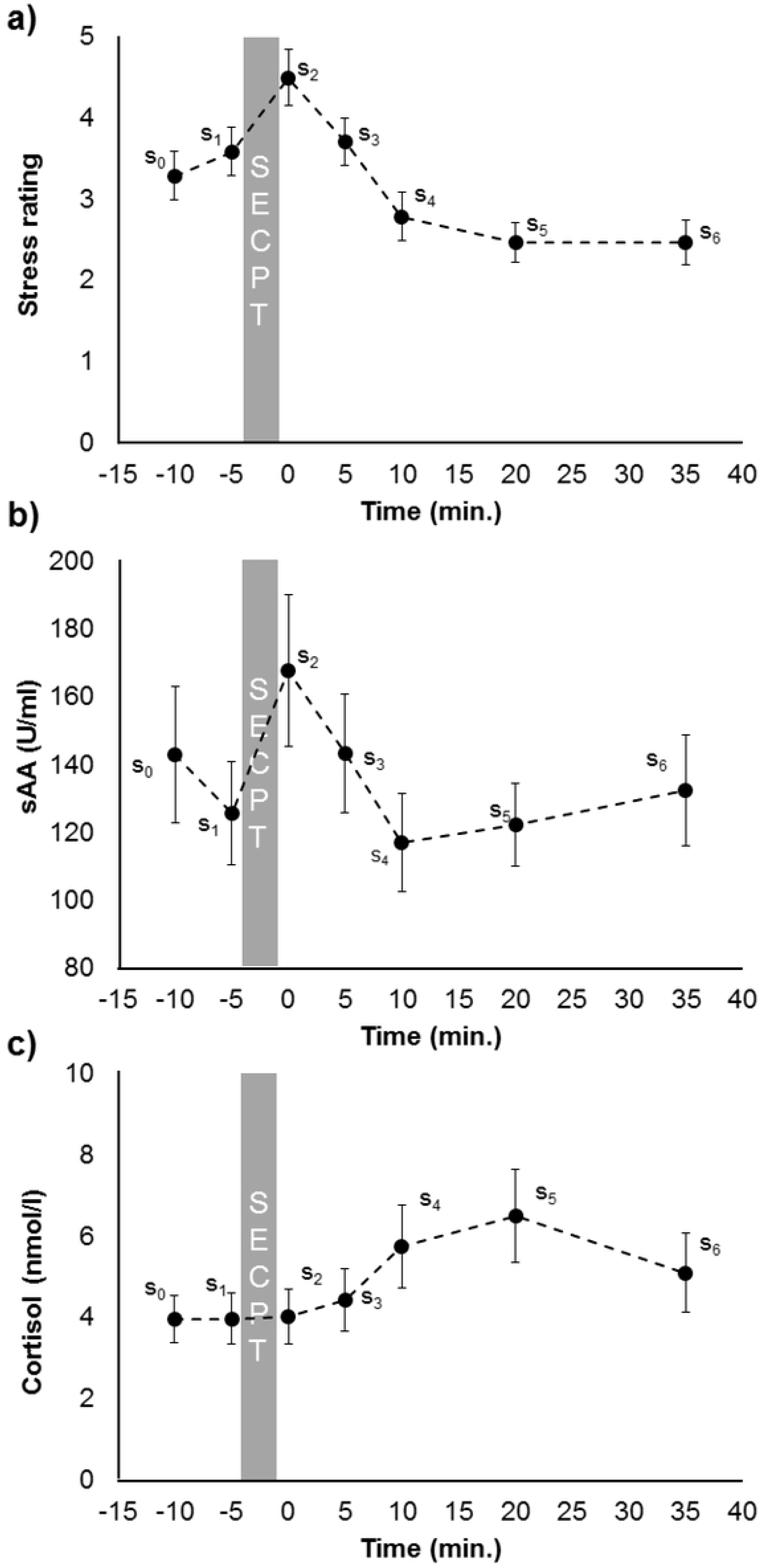
Time course of subjective stress ratings (a), sAA concentration (b), and cortisol concentration (c) at different time points before and after the SECPT.

### Salivary alpha-amylase

Salivary alpha-amylase concentration significantly differed between the seven time points (*F*_(6, 192)_ = 2.32, *p* = .034, η_p_^2^ = .07, Fig 4b). Post-hoc t-tests revealed that sAA concentration increased immediately after the SECPT (s_1_-s_2_: *t*_(32)_ = −2.20, *p* = .035, *d* = 0.49) and then decreased until it reached a minimum ten minutes after the SECPT (s_3_-s_4_: *t*(_32_) = 1.91, *p* = .033, *d* = −0.32). Amylase concentration did not change further after s_4_.

Twenty-two participants were classified as sAA-responders with an sAA-increase of more than ten percent between s1 and s2. Eleven participants were classified as sAA non-responders. A further rmANOVA with the within-subject factors memory type and time and the between-subject factor sAA respondence was calculated. This revealed a main effect of memory type (*F*_(1,31)_ = 24.25, *p* < .001, η_p_^2^ = .44), a main effect of time (*F*_(2, 62)_ = 4.49, *p* = .015, η_p_^2^ = .13), a memory type * sAA respondence interaction (*F*_(1, 1)_ = 5.29, *p* = .028, η_p_^2^ = .15), and a memory type * time interaction (*F*_(2, 62)_ = 15.41, *p* < .001, η_p_^2^ = .33).

Post-hoc analysis for the PE revealed only a main effect of time (*F*_(2, 62)_ = 7.55, *p* = .001, η_p_^2^ = .20; Fig 5a). Thus, the PE and, therefore, long-term memory performance was not associated with the sAA response. For the RE, a main effect of time was found (*F*_(2, 62)_ = 18.24, *p* < .001, η_p_^2^ = .37) and a main effect of sAA respondence were found (*F*_(1, 31)_ = 4.77, *p* = .037, η_p_^2^ = .13; Fig 5b). Only the sAA responders showed a main effect of the factor time effect (*F*_(2, 42)_ = 23.37, *p* < .001, η_p_^2^ = .53), but not the sAA non-responders. Therefore, WM performance only increased in sAA responders.

**Fig 5.**
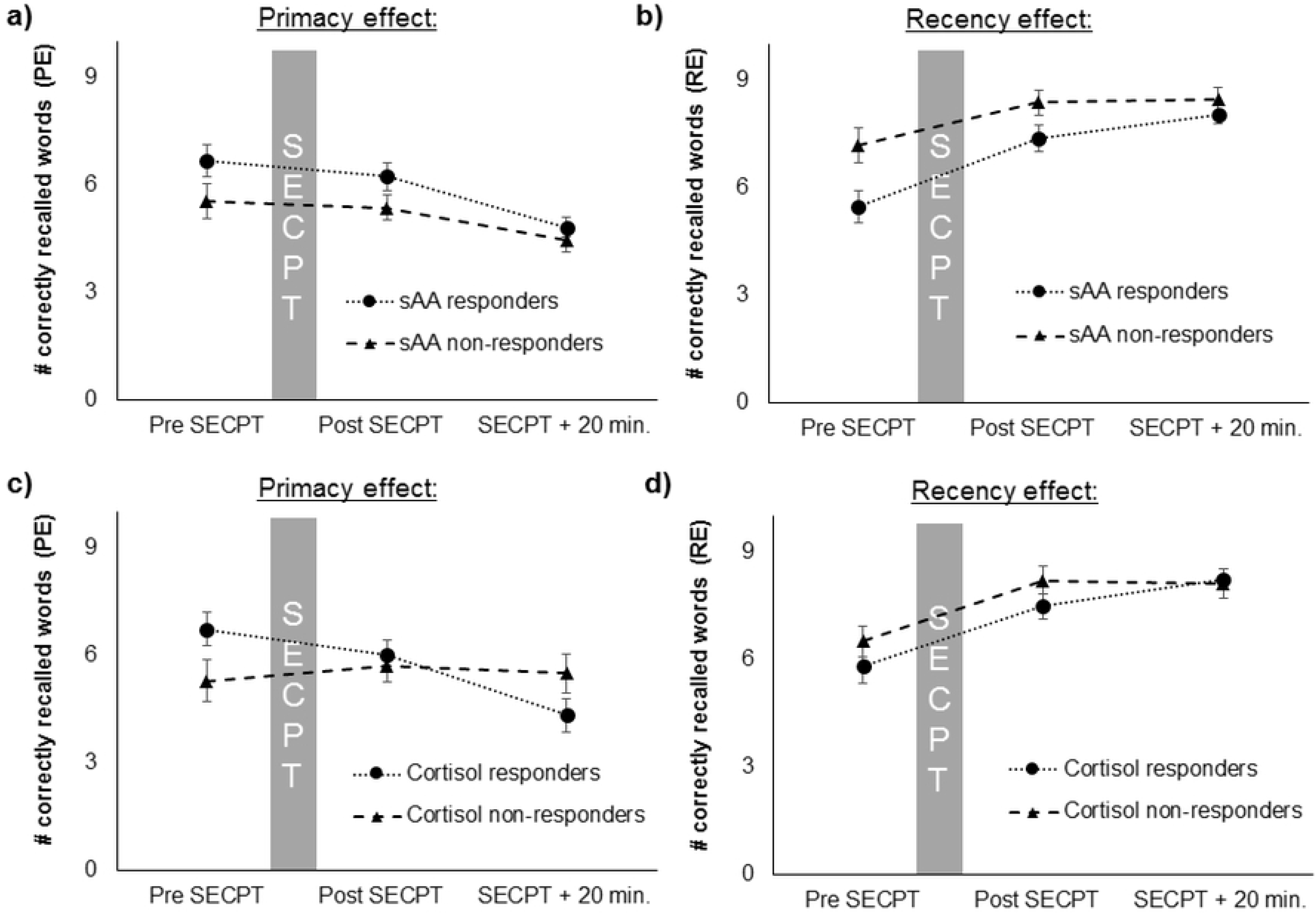
Primacy and recency effects for sAA (a, b) as well as for cortisol (c, d) responders and non-responders, before, after and 20 minutes after the SECPT.

### Cortisol

Cortisol concentration also differed significantly between the seven time points (*F*_(2.02, 164.48)_ = 7.08, *p* = .001, η_p_^2^ = .20, Fig 4c). Post-hoc t-tests revealed that cortisol concentration did not differ between before and immediately after the SECPT (*p* = .368). Afterwards cortisol concentration increased until it reached a maximum 20 minutes after the SECPT (s_4_-s_5_: *t*_(32)_ = – 3.35, *p* = .002, *d* = 0.45).

Twenty-three participants were classified as cortisol-responders with a cortisol increase of more than ten percent between s_1_ and s_5_. Ten participants were assigned to the cortisol non-responders group. A further rmANOVA with the within-subject factors memory type and time and the between-subject factor cortisol respondence was calculated. This revealed a main effect of memory type (*F*_(1, 31)_ = 16.51, *p* < .001, η_p_^2^ = .35), a main effect of time (*F*_(2, 62)_ = 7.01*,p* = .002, η_p_^2^ = .18), a memory type * time interaction (*F*_(2, 62)_ = 13.12, *p* < .001, η_p_^2^ = .30), and a memory type * time * cortisol respondence interaction (*F*_(2, 1)_ = 4.07, *p* = .022, η_p_^2^ = .12).

Post-hoc analysis for the PE revealed a main effect of time (*F*_(2, 62)_ = 4.92, *p* = .01, η_p_^2^ = .14) and a time * cortisol respondence interaction (*F*_(2, 31)_ = 11.79, *p* = .005, η_p_^2^ = .16). Post-hoc analyses showed that, for the PE, a time effect was found only for the cortisol responders (*F*_(2, 44)_ = 18.28, *p* < .001, η_p_^2^ = .45), but not for the cortisol non-responders (Fig 5c). Therefore, long-term memory performance only decreased in cortisol responders after the SECPT, but not in cortisol non-responders. For the RE, only a main effect of time (*F*_(2, 62)_ = 15.44, *p* < .001, η_p_^2^ = .51), but no interaction time * cortisol respondence or a main effect of cortisol respondence were found (Fig 5d).

## Discussion

In this study, we investigated the time course of the physiological stress response and its associations with WM and LTM performance. The latter were operationalized by means of the PE and the RE of the serial position curve. Our first finding was that WM performance increased immediately after the stressor in participants who showed a sAA response. No changes in WM performance were found in sAA non-responders. This is in line with our hypothesis and with previous findings, in which improvements in WM performance after an acute stressor were found as well (38–42). However, other studies also found the opposite, i.e. impaired WM functioning after an acute stressor (e.g., 43–47). However, in contrast to our study, in most cases spatial WM and not verbal WM was investigated in these previous studies. One explanation that has been proposed to explain the different findings was that WM improves for simple tasks. but that it is impaired for complex tasks (2). This explanation fits well to the results of our study because we used simple word lists with only 15 non-emotional words, and it can be assumed that this was an easy task for our participants.

Our second finding was that LTM performance did not change immediately after the stressor, but decreased 25 minutes after its onset in cortisol responders. This was also the time point of the maximal cortisol response. This decrease in LTM performance was not found in cortisol non-responders. Our finding is in line with many previous studies in which a drop in LTM performance and a relation with glucocorticoids was found (e.g., 48; 49–52). However, there are also a few studies which cannot support this conclusion (53, 54). It has been proposed that the timing of the glucocorticoid release (or injection for pharmacological studies), is the critical factor for the diversity of the findings (55, 56, 56, 57). This could also explain why we did not find an effect on LTM performance immediately after the stressor.

It is important to point out that neutral words were used in our study. There is an extensive literature on the effects of (e.g., emotional) arousal or stress on emotional memory (e.g., 58, 59). It was found that emotional LTM is enhanced – and not impaired as it is for neutral stimuli (60). For memory formation of emotional stimuli, the amygdala plays a critical role (61). Emotional memory is indeed affected by SNS activity (62). For example, it has been found that the enhancement in emotional LTM is eliminated through a blockade of beta-adrenergic receptors in humans (58). Furthermore, it was shown that a noradrenaline injection after learning enhances LTM for emotional stimuli (59). In animal studies, it has been found that noradrenaline can have long-term effects on the hippocampus (63–66). Besides, glucocorticoids are also involved in LTM enhancement for emotional stimuli. For example, (67) found that a cortisol injection during learning enhanced recall one week later. To combine these manifold findings, it has been suggested that both, the glucocorticoid and the noradrenaline pathway, interact in emotional memory formation (68, 69).

Furthermore, it should be noted that, in previous studies, LTM was assessed in a different way than in our study. In most previous paradigms, participants were presented the material on one day and the recall took place on another day. Thus, the elapsed time was much longer than in our study. Our design has the advantage that learning as well as retrieval of the items could both be tested at all three time points within one person. Therefore, our design offers new insights in the effects of acute stress on memory performance.

In future research, our design should be repeated with emotional stimuli. Furthermore, it should be combined with imaging techniques to get more insight into the underlying neural processes. Besides, our design should be supplemented by the collection of blood samples, because the use of sAA as marker for SNS activity is well-established (70, 71), there are still some valid concerns that need to be taken into account (72). Furthermore, our design should be supplemented by pharmacological treatments, which block either the MR or the GR, because to understand the underlying mechanisms both receptor types should be taken into account (16, 73).

## 5 Conclusions

Our study supports the assumption that SNS activation after an acute stressor immediately improves WM function. This is probably related with noradrenergic and dopaminergic activation of the PFC. Furthermore, we showed that HPA axis activity is associated with LTM processes – probably through interactions with the hippocampus. Using the serial position effects to measure both WM and LTM performance within one test seems to be a very good means for further research on the effects of acute stress on memory.

## Acknowledgement

We thank Aylin Gögsen, Kristin von Majewski, and Yvonne Daichendt for data collection.

## Supporting information

**S1 File. S1_Data.csv**. Data file.

